# Relative distance change rather than absolute distance drives auditory alerting

**DOI:** 10.64898/2026.11.25.769175

**Authors:** Petra Kovács, Brigitta Tóth, István Winkler, Francesco Pavani, Robert Baumgartner

## Abstract

The function of hearing as an early warning system relies partly on detecting the distance and motion of an object based on sound. Approaching — or looming — objects activate the human alerting system, but it is debated whether sounds in the immediate vicinity of the body achieve the same irrespective of the direction of motion and of sound intensity. We addressed this question by examining distance perception of and phasic alertness states evoked by virtual sonic motion into and within peripersonal and extrapersonal space, at high and low intensities, in normal-hearing adults. Based on distance ratings, target detection promptness, and electroencephalography (EEG), we found that looming auditory objects increase phasic alertness and auditory sensory gain, and we also observed a general tendency to perceive sound offsets as closer to the body than onsets. However, we did not find evidence that auditory objects moving at a constant distance within peripersonal space affect alerting over and above the influence of intensity. We suggest that the auditory looming bias is not amplified further by preferential processing of auditory information near the body, and that the early warning function of hearing emphasizes the direction of motion over distance.

**Significance statement:** Hearing acts as an early warning system, helping us sense and react to objects moving toward us even when we cannot see them. A long-standing debate is whether the brain reacts more strongly simply to sounds close to the body, or specifically to sounds that are approaching. Using moving sounds together with brain recordings (EEG) and reaction-time measures, we found that only approaching sounds heightened alertness and boosted the brain’s auditory response, while sounds circling at a fixed close distance did not. Listeners also tended to hear moving sounds as ending nearer than they began. These findings show that it is the change in distance, not nearness itself, that triggers the auditory alerting response underlying adaptive, defensive actions.

## 1. Introduction

Detecting the motion and distance of nearby objects ensures safety by helping to select suitable defensive actions – for example, fight or flight –, and hearing is particularly suited to this function for objects out of sight. Motion towards the body – i.e., looming – has especially high behavioral relevance. Compared to receding sources, looming sources are perceived as closer (Neuhoff et al., 2012; Rosenblum et al., 1993), faster (Neuhoff, 2016), and changing more in intensity (Neuhoff, 1998) in a phenomenon known as the auditory looming bias (ALB). Looming sounds also increase phasic alertness (Bach et al., 2008, 2009, 2015; Cappe et al., 2012), a component of attention enabling prompt reactions on a transient timescale (Petersen & Posner, 2012), and elicit distinct neural responses compared to receding sources (Bach et al., 2008, 2015; Cappe et al., 2012; Greif et al., 2025; Seifritz et al., 2002), which possibly reflects increased sensory gain, i.e., a response amplification of neurons processing especially relevant or informative inputs (Hillyard et al., 1998; Mehrpour et al., 2020). The dominant acoustic cue for looming perception is rising intensity (Ignatiadis et al., 2024; Rosenblum et al., 1987). However, spatial cues also contribute to alerting effects (Bach et al., 2009), and the ALB can be elicited by spectral cues alone (Baumgartner et al., 2017), especially if the sound is localizable in the listener’s surroundings (Greif et al., 2025).

Sounds near the body are behaviorally relevant even when they are not looming. The space within which one can interact with objects, termed peripersonal space (PPS), is characterized by increased multisensory integration, which helps predicting proximal events based on distal input and plan suitable motor action (Graziano et al., 1999; Ladavas et al., 2001; Serino et al., 2009; Zampini et al., 2007; Zanini et al., 2021). Auditory distance relative to the body can be determined not only based on intensity, but also through other monaural and binaural auditory cues (for a review see Kolarik et al., 2016). However, auditory distance perception is imprecise: near sources are overestimated, while far sources are underestimated (Zahorik & Wightman, 2001), making the border between auditory PPS and extrapersonal space (EPS) fuzzy.

It remains unclear whether near sources increase phasic alertness and sensory gain independently of looming motion. The ALB takes effect even if the looming source stays in EPS (e.g. Neuhoff, 2001), but some studies suggest that phasic alertness and motor preparation scale together with how close the source looms into PPS (Bahadori et al., 2021; Camponogara et al., 2015), suggesting a possible unisensory, auditory bias towards PPS. However, these effects were specific to looming sounds and absent for receding or stationary controls, suggesting a lack of a PPS bias independently of ALB. In contrast, other studies reported higher motor excitability or multisensory integration in PPS regardless of motion direction (Canzoneri et al., 2012; Finisguerra et al., 2015).

Finally, interpretation is confounded by the effects of temporal predictability and sound intensity. Sounds that loom closer into PPS frequently coincide with longer trials; therefore, reaction times may reflect increasing response likelihood as a trial unfolds besides a true effect of the ALB. Moreover, responses to sounds in PPS are confounded with responses to high sound intensity in general, even though near sources and loud sources are not always equivalent (Brungart et al., 1999; Kopčo et al., 2012; Kopčo & Shinn-Cunningham, 2011; Little et al., 1992).

Overall, conflicting results and confounds between temporal predictability, distance, and intensity make it unclear whether auditory alerting and sensory gain is driven by looming motion, proximity to the body, or both. In this preregistered study (Kovács et al., 2025), we examined this question hypothesizing that phasic alertness and sensory gain increase in response to sounds that either loom towards the listener (H1) or move within their PPS (H2). We collected distance ratings as well as behavioral and electroencephalographic (EEG) data for binaurally rendered sounds moving in radial or angular trajectories and ending near, in PPS, or far, in EPS.

## 2. Methods

### 2.1. Participants

Twenty-six young adults participated in the study (15 female, 11 male, mean age 24.23 years, SD = 3.42, 25 right-handed). Four participants were excluded due to data collection error (malfunction of the earphones) or low EEG data quality (see Section 2.4. EEG data acquisition and preprocessing); therefore, the final dataset includes 22 participants (12 female, 10 male, mean age 24.36 years, SD = 3.50, 21 right-handed). Additionally, participants were reinvited for a second occasion to perform an auditory distance rating task. Eleven participants returned for this task (5 female, 6 male, mean age 26.45 years, SD = 2.97).

None of the participants had indications of hearing disorders as confirmed by pure tone audiometry at six octave frequencies between and including 250–8000 Hz: <30 dB hearing thresholds, <20 dB difference between ears. No participant had a history of neurological disorders or used any medication affecting the nervous system.

The study fully complied with the World Medical Association Helsinki Declaration, all applicable national laws, and the ethics statement for human experimental research conducted at the Acoustics Research Institute of the Austrian Academy of Sciences, as approved by the Commission for Science Ethics of the Austrian Academy of Sciences on April 23rd, 2024. Before the experiment began, participants were informed of their rights — including the right to withdraw at any time without penalty — and provided informed consent. Instructions were given both orally and in writing, and participant well-being was closely monitored throughout the procedures. All participants received financial compensation for taking part.

### 2.2. Stimuli

The stimuli were cue-target sound pairs presented over earphones (Figure 1A). There were 2✕2✕2 stimuli conditions based on cue motion trajectory (radial or angular), motion offset distance (offset in EPS, offset in PPS) and source intensity (high, low). Note that radial sounds moving into PPS can be identified as looming, whereas radial sounds moving into EPS can be identified as receding. The characteristics of the control stimuli in studies of the PPS are worth mentioning. Previous studies used either stationary (e.g. Camponogara et al., 2015) or receding (e.g. Canzoneri et al., 2012; Finisguerra et al., 2015) controls. The advantage of the former is that distance is kept constant, enabling the investigation of its unique role independently of motion. The advantage of the latter is that moving sound sources are compared to other moving sound sources. Our approach uniquely combined both advantages by using rotating controls at different distances.

**Figure 1.**
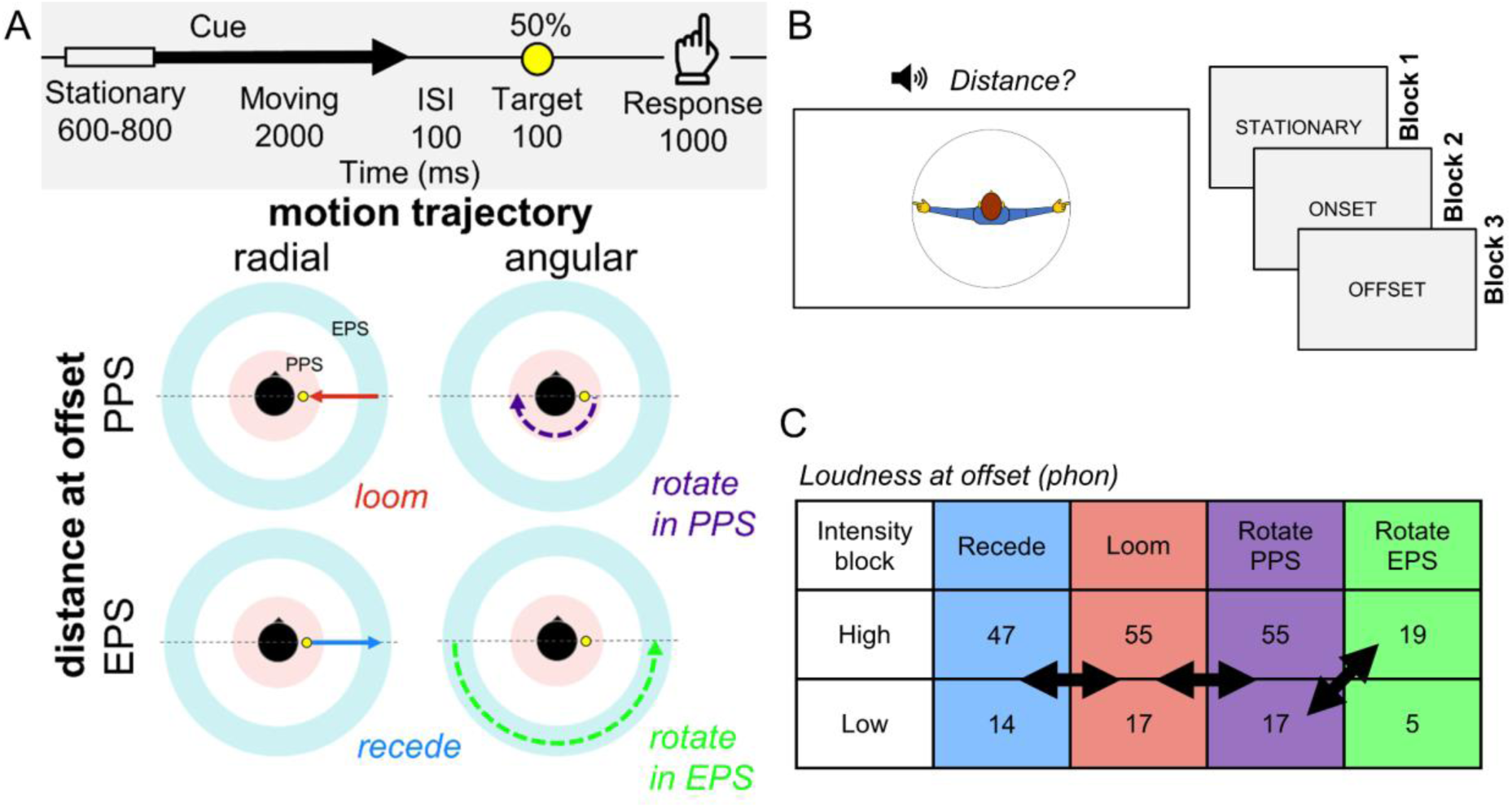
Study design. **A**: Design of the main task. Upper panel: the timeline of a trial; lower panel: cue conditions (the yellow circle indicates the position of the target, always shown at +90° for illustrative purposes). **B:** Design of the distance rating task. Left panel: the graphical user interface presented to the participants; right panel: order of blocks. Each block required a rating about the perceived position of a stationary sound source, of the onset or the offset of a moving sound source, respectively. **C:** Average loudness of sounds at offset in each cue condition and intensity block, assessed according to the binaural loudness model from Moore et al. (2018). Arrows denote the conditions contrasted in EEG data analysis.

The goal of having source intensity as an independent variable was to enable its dissociation from source distance. Therefore, sounds in PPS in the low intensity condition had the same absolute intensity as sounds in EPS in the high intensity condition, while the intensity range in the two conditions was identical (i.e., the relative intensity was informative of distance; Figure 1C). To achieve this, the intensity of sound sources was made 20 dB lower in the low intensity condition, which corresponds to the difference in sound pressure level (SPL) between sounds at 0.2 m and 2 m in the free field (although note that our specific stimuli were simulated in a reverberant environment, and therefore individual looming and receding sounds encompassed a smaller SPL range). The SPL of high intensity sounds ranged from 60-80 dB, low intensity sounds from 40-60 dB as measured by a Brüel & Kjær 2260 sound level meter.

Cue sounds (Figure 1A) were triangle waves, as previous studies reported larger perceptual salience for looms of harmonic sounds (associated with single sources such as an animal) than of noise (associated with diffuse sources like the wind; Ghazanfar et al., 2002; Neuhoff, 1998, 2001). The fundamental frequency was varied in one octave range (310-620 Hz, linearly spaced unique values) to avoid auditory habituation effects as much as possible for cleaner ERP measurements. Cues had a duration of 2600-2800 ms with 100 ms on– and offset ramps. They each started with a stationary portion whose duration was randomized between 600-800 ms for less predictable motion onset. This was followed by a 2000 ms moving portion. Including a stationary onset allows for dissociating ERP responses to *motion* onset from *sound* onset (Baumgartner et al., 2017; Getzmann & Lewald, 2010) — we only analyzed motion ERP responses. Cues were spatialized at predefined distances based on our pilot results evaluating the perception of auditory PPS and EPS. Looming and Receding sounds moved between 0.2 m (PPS) and 2 m (EPS). Sounds rotating in PPS moved at a constant distance of 0.2 m. Sounds rotating in EPS moved at a 20 m distance to ensure comparable levels between rotating in EPS, high intensity and rotating in PPS, low intensity conditions. Looming and receding cues had a fixed azimuth of +90° or –90° (left or right side, randomized), while rotating cues moved from +90° to –90° azimuth or vice versa (left to right or right to left, randomized) via the listener’s back (staying outside the listener’s field of view). Sounds moved at a constant speed of 0.9 m/s for radial motion and 90°/s for angular motion. All sounds had a constant elevation of 0°.

Target sounds were brief sinusoid tone pips following the methods of Bach and colleagues (2009). They had a frequency of 2400 Hz and a duration of 100 ms with 10 ms on– and offset ramps. Every target was spatialized in PPS (at 0.2 m), with an azimuth of +90° or – 90° (randomized) and an elevation of 0°. They were presented after 100 ms inter-stimulus-interval (ISI) following the cue. Targets appeared on only half of the trials. Half of the targets were presented on the same side as the preceding cue offset (congruent targets), while half were presented on the opposite side (incongruent targets).

Both cues and targets were presented binaurally. The cues were spatialized with the BeRTA Renderer Application v3.6.0 for Windows x64, using BRT v2.2.0 (González-Toledo et al., 2023) controlled by Open Sound Control through custom Matlab scripts. The motion percept was supported by spectral, intensity, and reverberation cues. A KEMAR HRTF from the SADIE II database (Armstrong et al., 2018), measured at 1.2 m, was used for filtering. A scattering delay network environment model was enabled for spatialization, using both the direct and the reverberation path. Near-field correction of ILDs was implemented as a difference filter, which models the spectral changes between a source at a far-field measurement distance and a simulated near-field distance, at identical azimuth and elevation (Cuevas-Rodríguez et al., 2019). Cross-ear parallax correction was also applied, which accounts for the modified angle of incidence at each ear, compared to the angle of incidence measured at the center of the head (Cuevas-Rodríguez et al., 2019). Barycentric interpolation was enabled based on the three nearest HRTF positions to the simulated sound position. Distance attenuation was applied so that doubling the source-listener distance resulted in a 3 dB sound level reduction. The targets were spatialized on a different platform than the cues because they were too brief to be handled by BeRTA through Matlab with precise timing. Therefore, the SOFA toolbox v.2.1 (Majdak et al., 2022) was employed using the same HRTF set as for the cues but without any room simulations and distance adjustments. The sampling rate of both cues and targets was 48 kHz. There were 100 stimuli repetitions per cue condition, separately for each intensity condition, resulting in 400 trials per intensity condition, altogether 800 trials.

### 2.3. Procedure

#### Target detection task

Participants sat in a soundproof room and were presented with the sound stimuli through insert earphones (ER-2, Etymotic Research) on an RME Fireface UC audio interface. The experimenter explained the task to the participant and presented them with example stimuli both with and without a target.

During the task, participants were instructed to listen to each sound stimulus and detect any targets as quickly and accurately as possible with a keypress. EEG and skin conductance response (SCR) data were recorded during the main task phase. Stimuli were presented in 10 blocks of 80 trials, each, with optional short breaks between the blocks. The first 5 blocks contained either only high– or only low-intensity sounds (counterbalanced across participants), while the second 5 blocks were dedicated to the other source intensity condition. An obligatory short break followed after the first 5 blocks. Each trial consisted of a cue (2600–2800 ms), an ISI (100 ms), and either a target or silence (100 ms), adding up to 2800–3000 ms per trial (Figure 1A). The inter-trial interval (ITI) was randomized between 2600–2800 ms, with round 100 values (for less predictable timing and therefore cleaner ERPs). The response window for keypresses was 100–1000 ms from target onset. The whole experiment took ∼3.5 h including audiometry and the setup of EEG and SCR hardware.

In each stimulus block, each of the four cue motion trajectories (i.e., looming, receding, rotating in PPS, rotating in EPS: Radial vs. Angular ✕ PPS vs. EPS) received a quarter of the trials (20 trials per block), presented in randomized order. All order randomizations occurred independently of the previous trials without constraints. Each of the four trial types received 10 target and 10 non-target trials within each stimulus block, and the order between them was randomized. The cue onset azimuths (+90° and –90°) were also presented in random order: their number was balanced across the 5 blocks, but not within each block, separately. The number of congruent and incongruent target trials was balanced within each intensity condition, but not within each block. Randomizations were done separately for each participant. Participants received feedback about their accuracy and reaction time (RT) after each stimulus block.

Stimulus presentation was controlled from PsychToolbox (v3.0.19) running on Matlab 2023b. Participants were instructed to press the SPACE key with their right hand on a computer keyboard in response to target sounds, and the accuracy and promptness of each response was recorded. A computer screen was used to show the instructions as well as a fixation cross during the task. Participants rested their chin on a chinrest to minimize head movements and were instructed to keep their eyes on the fixation cross during sound presentation.

#### Distance rating task

Initially, the distance rating task completed by all 22 participants delivered separately generated sound stimuli, which were not exactly identical to the ones presented in the main task. The stimuli were generated separately, with no stationary onset, although otherwise according to the same pipeline as described in the Stimuli section. The task was completed separately for two source intensities (high, low) with the order identical to the order of source intensities in the main task (and thus counterbalanced across participants). Participants completed the distance rating task at a given intensity, then the target detection task at the same intensity, followed by another round of the distance rating and target detection task at the other intensity. Due to the unintended inconsistency of stimuli across the two types of tasks, we decided to exclude this data from analysis. Instead, we repeated the distance rating task with the same stimuli as was used in the main task by reinviting the participants. We only report here the results collected during the reinvitation session.

Distance ratings were collected for both stationary and moving sounds (Figure 2B). The moving sounds were chosen from the stimulus material of the target detection task (only the cues, without targets). 20 stimuli – 10 of each offset azimuth – were chosen from each condition (looming, receding, rotating in PPS, rotating in EPS). While the duration of the stationary onset and the F0 were counterbalanced in the stimulus material of the target detection task, the stimuli used in the distance rating task were picked with the intention of counterbalancing only cue condition and offset azimuth, involving random combinations of stationary onset duration and F0 to reduce the number of ratings that participants had to provide. For the 80 moving sounds (4 cue conditions ✕ 2 azimuths), participants were instructed to judge the position (see below) of the sound source at sound onset in the first block of distance-judgement trials for moving sounds, then the position at sound offset in the next block of trials in fixed order.

**Figure 2.**
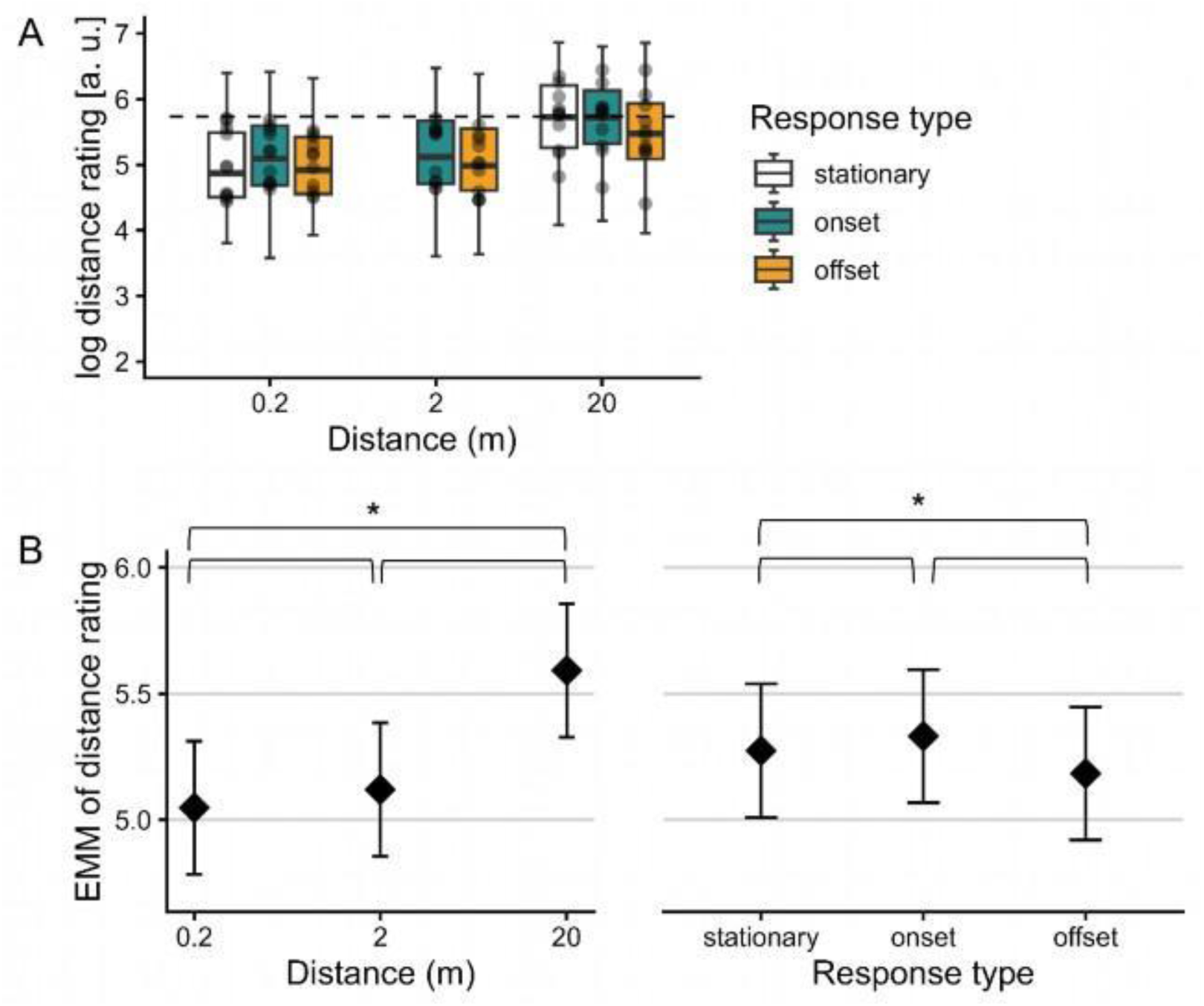
Results of the distance rating task. **A**: Summary of log-transformed distance ratings for stationary and moving sound sources. The dashed horizontal line denotes the boundary of PPS (shown as a circle in Figure 1B). Points show average responses per participant. Note that two intermediate distances tested for stationary sources only (3.2 and 9.8 m) are not shown in the figure. **B:** Estimated marginal means (EMM) of log-transformed distance ratings as a function of distance (averaged over all response types, left panel) and by response type (averaged over all distances, right panel). Asterisks denote a significant difference. Error bars: 95% CI.

An additional 80 stationary sounds were created according to the cue spatialization pipeline described in Section 2.2. Stimuli. They were binaurally rendered at the distances used for PPS (0.2 m) and EPS (20 m) stimuli, as well as two intermediate distances (3.2 and 9.8 m). Azimuths of +90° or –90° were used throughout the whole 2 s sound duration. For the 80 stationary sounds (20 for each distance), participants had to judge the perceived position (see below) of the sound source.

Participants were instructed to listen to each sound as many times as they wished and estimate the position of the sound source with a mouse click on a graphical user interface (Figure 1B), especially noting whether the sound originated from inside or outside arm’s reach. The user interface depicted a person from a bird’s eye view with arms spread to the side, and the distance of the arm’s reach was marked by a circle. To encourage an externalized percept, the participants were told to think of the sounds as originating from a fly and estimate its position. Upon mouse click, an icon of a fly appeared on the screen in the corresponding position. The participants controlled the sound playback by clicking on a loudspeaker symbol on the screen. They were able to both listen to the stimulus and change their rating as many times as they wished within the trial. After they were satisfied with their rating, they progressed to the next trial with a button press and could not navigate back to previous trials again.

A low– and a high-intensity run was administered to the participants in the same order they had encountered these intensities in the main task, with a short break between them. A total of 480 ratings were given by each participant, 240 in each intensity run (80 stationary, 80 onset, and 80 offset ratings, always in this order; Figure 1B) in response to a total of 160 unique sounds (80 stationary and 80 moving). The task lasted about 1 h.

### 2.4. EEG data acquisition and preprocessing

EEG data were acquired with an actiCHamp (Brain Products) system with ActiCap active Ag/AgCl electrodes. There were 64 electrodes, including the reference electrode FCz, applied according to the international 10/20 system. Four of the 64 electrodes (FT9, FT10, TP9, TP10) were placed around the eyes to monitor ocular movements. Conductive gel was used as conductive medium. All electrode impedances were kept below 25 kΩ at the beginning of data collection. EEG was recorded at 1 kHz sampling rate with 100 Hz low-pass filtering. Audio triggers were passed through a StimTrak device and recorded as auxiliary input to correct audio playback latencies later on. BrainVision Recorder software was used for EEG recording.

EEG preprocessing followed in EEGLAB 2024.2 (Delorme & Makeig, 2004) running on Matlab 2023b (The MathWorks Inc, 2023). First, the EEG data were imported into EEGLAB, then latency corrections were performed on the EEG triggers based on the StimTrak audio triggers. EEG and SCR data were separated at this point. EEG data were band-pass filtered between 0.1–30 Hz, then re-referenced to the common average, keeping the original reference channel (after adding an extra channel of zeros). After this, extreme artifacts were rejected based on visual inspection of the data; specifically, this included rejecting breaks between experimental blocks, extreme motion artifacts and noisy channels. A maximum of 6 channels (10% of all electrodes) were rejected for each participant. Then, independent component analysis (ICA) decomposition was run using the extended InfoMax algorithm. Blink and eye-movement artifacts were then rejected based on visual inspection of the topographical distribution, as well as temporal and spectral characteristics of the components. Maximally six components (10%) were rejected. The data quality of two participants was too low to detect distinct ICA components; therefore, their data were excluded from further analysis (as mentioned in Section 2.1. Participants). Previously removed channels were then interpolated using spherical interpolation. Data were epoched between –200 and 1000 ms from sound motion onset, separately for each motion trajectory ✕ offset distance ✕ source intensity combination. Data were downsampled to 100 Hz. Epochs containing a >100 µV amplitude change were rejected from further analysis. The average number of rejected EEG epochs in the final dataset was 21 for the looming and rotating in PPS, and 20 for the receding and rotating in EPS conditions, out of a total of 200 per condition.

### 2.5. Skin conductance data acquisition and preprocessing

Skin conductance data were acquired with a galvanic skin response (GSR) sensor (Brain Products) integrated with the EEG system. Two electrodes were used, one on the index and one on the middle finger (middle phalanx) of the left hand. Data were recorded at 1 kHz sampling rate with 100 Hz low-pass filtering. BrainVision Recorder was used for data recording.

The data were preprocessed in EEGLAB 2024.2 running on Matlab 2023b. Latency-corrected SCR data were bandpass filtered between 0.1–5 Hz (Bach et al., 2009), then epoched between –1000 and 5000 ms from sound motion onset, separately for each motion trajectory ✕ offset distance ✕ source intensity combination, using only no-target trials, which did not require a button press response from the participants. Epoched data were downsampled to 100 Hz. All 400 SCR epochs per participant were used in the analysis.

### 2.6. Statistical analysis

#### 2.6.1. Preregistered analyses: Behavioral data

Target detection promptness (1/RT) data were analyzed in R (R Core Team, 2024) using the ‘lme4’ (Bates et al., 2015) and ‘emmeans’ (Lenth, 2023) packages. Promptness was recorded for each target trial and averaged across participants, separately for each motion trajectory ✕ offset distance ✕ source intensity combination. Inaccurate button press responses (false alarms), as well as responses with an RT outside of the 100–1000 ms time window from target onset were excluded from analysis. A linear mixed-effects model was used to analyze promptness data. Source intensity was converted from categorical (high vs. low) to a continuous value by calculating the intensity of each cue stimulus at the offset, averaging over the last 100 ms of the louder channel excluding the offset ramp, and was expressed in dB full-scale (dB FS). In the linear mixed model, offset intensity, motion trajectory (radial v. angular), and offset distance (PPS v. EPS) were treated as fixed effects, and participant as a random effect (random intercept). The model additionally included an interaction term between motion direction and offset distance.

#### 2.6.2. Preregistered analyses: EEG data

The preprocessed EEG data were converted to FieldTrip format (Oostenveld et al., 2011). The –200–0 ms baseline was removed and average ERPs to sound motion onset were calculated for each participant and condition in FieldTrip. The ERP data were then converted to ‘.fiff’ format in order to perform the interim analysis in MNE Python (Gramfort et al., 2013).

The interim analyses were performed in the ‘niseq’ package for MNE Python (Veillette et al., 2023). The first look at the data occurred after the 10th participant’s data had been collected, and the second look after the 22nd data collection. The purpose of the interim analyses was to test H2 (phasic alertness is higher in PPS than EPS). For this purpose, difference waveforms were calculated for ERPs evoked by angular cues with EPS offset minus angular cues with PPS offset, pooling together the two intensity conditions. A one-sample, two-tailed cluster-based permutation analysis was performed on the difference waveform with the maximum number of participants for the sequential analysis set to 30, the alpha level set to 0.05, with 50 permutations and a linear alpha spending function.

The comparison of ERPs evoked by the radial cues (related to H1, i.e., higher phasic alertness for looming than receding cues, and thus not included in the interim analysis) were only analyzed once the full dataset was collected. Note that this also holds for all behavioral and SCR data (not included in the interim analysis either). ERPs evoked by radial cues were baseline-corrected and averaged in FieldTrip, then converted to ‘.fiff’ format and analyzed in MNE Python. Difference waveforms were calculated between ERPs evoked by radial cues with EPS offset (receding) and radial cues with PPS offset (looming). The difference waveform was analyzed with a one-sample, two-tailed cluster-based permutation test with 500 permutations.

#### 2.6.3. Preregistered analyses: SCR data

The epoched SCR data were baseline-corrected (–1000–0 ms) in FieldTrip (Oostenveld et al., 2011). The average SCRs evoked by the cues were calculated separately for each motion trajectory x offset distance condition. These were then analyzed in MNE Python using a cluster-based permutation test with 1000 permutations.

#### 2.6.4. Exploratory analyses: Behavioral data

Besides the preregistered analyses (Kovács et al., 2025), exploratory analyses were also performed on the behavioral data. Firstly, we analyzed the results of the distance rating task in R using the ‘lme4’ and ‘emmeans’ packages. The aim was to compare distance ratings as a function of response type (stationary, onset, and offset responses) and distance (PPS and EPS m). Distance ratings provided for the stationary sounds at the two intermediate distances (3.2 and 9.8 m) were excluded from this analysis. First, a linear mixed-effects model was fit on the log-transformed distance ratings, with response type and distance as fixed effects and participant as random effect. Then, the estimated marginal means were calculated separately for response type and distance by keeping the other fixed effect constant in both cases.

Secondly, we analyzed the effect of cue-target spatial congruence on promptness in the target detection task. We fit a linear mixed effects model where we treated motion condition as a fixed effect with 4 levels (looming, receding, rotating in PPS, rotating in EPS) and congruence as a fixed effect with 2 levels (Congruent, Incongruent), including their interaction. Participant was treated as a random effect. Note that in the previous analyses, motion condition was treated as 2 separate 2-level factors, as we aimed to avoid a 3-way interaction in this model.

Finally, we tested whether participants were able to dissociate intensity and distance by running a paired Student’s *t*-test on the log-transformed distance rating data comparing low-intensity sounds rotating in PPS with high-intensity sounds rotating in EPS. This analysis may elucidate the behavior behind the corresponding ERP comparison.

#### 2.6.5. Exploratory analyses: EEG data

ERPs obtained in the target detection task were explored within a longer time window than in the preregistered analysis, encompassing the whole length of the moving sound and the target using cluster-based permutation testing, with an additional cortical source estimation analysis. The same epoching procedure was used as described earlier (see Section 2.4. EEG data acquisition and preprocessing) but using a time window of –200 to 2200 ms from motion onset. Statistical analysis followed in MNE Python. Difference waveforms were calculated separately for the radial (looming minus receding) and the angular (rotating in PPS minus rotating in EPS) conditions, as well as for the conditions with PPS offset (looming minus rotating in PPS). The angular difference waveform was additionally tested in the trials where source intensities were comparable, i.e., rotating in EPS, high intensity subtracted from rotating in PPS, low intensity. Each of these difference waveforms was then subjected to a non-parametric cluster-level *t*-test with 1000 permutations.

The cortical sources of the difference waveforms were estimated in Brainstorm (Tadel et al., 2011) using the minimum-norm imaging method. Current density maps were normalized with the standardized low-resolution electromagnetic tomography (sLORETA) approach (Pascual-Marqui, 2002). The resulting current density maps were then tested against the 200 ms epoch baseline with a parametric Student’s *t-*test in order to explore statistically meaningful differences in source activation between conditions. False Discovery Rate (FDR) correction was applied to control for multiple comparisons, and the alpha level was set to *p* = 0.001 in this analysis.

Additionally, neurobehavioral models were fit on the promptness data to explore the connection between the EEG and behavioral results. The neurobehavioral models were based on the same linear mixed-effects model described in Section 2.6.1, except that they contained one additional fixed effect each: the EEG amplitude in the spatiotemporal (channel-time) cluster that was found significant in the exploratory cluster-based permutation analysis in the looming-receding comparison, the looming-rotating in PPS comparison, and the rotating in PPS-rotating in EPS comparison (see Section 3.4). To this end, ERP amplitudes were extracted in each trial containing a target (max. 400 trials per participant), taking the amplitude values within the significant spatiotemporal clusters and averaging them, separately for each participant and target-detection trial. Each cluster amplitude value was then paired with the observed target detection promptness in the corresponding trial. Trials missing from the EEG data (due to artifact removal) were also discarded from the behavioral data for this analysis. Finally, a behavioral and two neurobehavioral models (one per cluster contrast) were fit on these data, and their model fits were compared based on the Bayesian Information Criterion (BIC), where a smaller value means a more parsimonious and therefore better model.

### 2.7. Transparency and openness

This study was preregistered on OSF (Kovács et al., 2025). Experimental and analysis scripts are available in a GitHub repository (https://github.com/kovacs-petra/cherish). Anonymized raw EEG and SCR data, as well as behavioral data from all participants are available on a data repository (https://doi.org/10.60887/n2tamqt6r3pk8bk7).

## 3. Results

### 3.1. Behavioral results

#### 3.1.1. Distance rating task

The perceived distance of stationary sounds (simulated at 0.2, 3.2, 9.8, or 20 m), as well as the onset and offset of moving sounds (simulated at either 0.2, 2, or 20 m depending on the motion trajectory) was assessed in 11 reinvited participants. The goal of this exploratory analysis was to confirm the validity of the distances used in the main task. Figure 2 shows the results of this task in terms of distance ratings (panel A) and modeling results (panel B).

We first analyzed log-transformed distance ratings as a function of simulated distance by fitting a linear mixed effects model with response type (stationary, onset, offset) and distance (0.2 m, 2 m, 20 m) as predictors and then estimating the marginal means of the model. This analysis revealed two main results. Firstly, keeping response type constant, sounds at 0.2 m were rated at significantly smaller distances from the listener (*M_adj_ = 5.04*) than sounds at 2 m (*M*_adj_ = 5.13; *SE* = 0.02, *95% CI* [-0.12, –0.04], *p* < 0.001) as well as 20 m (*M*_adj_ = 5.62; *SE* = 0.02, *95% CI* [-0.61, –0.54], *p* < 0.001), while distance ratings of sounds at 2 m and 20 m were also significantly different from each other (*SE* = 0.02, *95% CI* [-0.54, –0.45], *p* < 0.001). This confirms that listeners successfully distinguished the selected distances (Figure 2B, left panel; see also Figure 2A).

Secondly and more unexpectedly, sound offsets were rated at significantly smaller distances (*M*_adj_ = 5.18) than both sound onsets (*M*_adj_ = 5.33; *SE* = 0.01, *95% CI* [-0.18, –0.11], *p* < 0.001) and stationary sounds (*M*_adj_ = 5.27; *SE* = 0.02, *95% CI* [-0.13, –0.04], *p* < 0.001), even if the effective distance was the same. Stationary sounds were also rated slightly but significantly closer than onsets (*SE* = 0.02, *95% CI* [-0.10, –0.01], *p* < 0.001). In other words, sounds were generally perceived as ending closer to the listener than where they started (Figure 2B, right panel; see also Figure 2A).

We further compared distance ratings provided for the onsets and offsets of moving sounds in the conditions where intensity was comparable but simulated distance was different. Specifically, log-transformed distance ratings for low-intensity sounds delivered in the rotating in PPS condition were compared to those for high-intensity sounds delivered in the rotating in EPS condition. High-intensity sounds rotating in EPS were rated at a significantly larger distance (*M* = 5.50) than low-intensity sounds rotating in PPS (*M* = 5.14; *t*(877.81) = 9.67, *p* < 0.001, *d* [95% CI] = 0.65 [0.79, 0.52]). Thus, listeners were able to dissociate sound intensity and distance.

#### 3.1.2. Target detection task

We examined how target detection promptness – as a measure of phasic alertness – was influenced by our experimental manipulations (Figure 3). Based on H1, we expected that promptness following looming sounds is larger than following receding sounds, whereas H2 predicted that any sound with a PPS offset would increase subsequent detection promptness compared to sounds with an EPS offset. The linear mixed-effects model fit on the target detection promptness data showed a significant effect of sound motion trajectory (angular vs. radial, *β* = –0.07; *95% CI* [-0.14, –0.004], *p* < 0.05), a significant effect of offset intensity (*β* = 0.005; *95% CI* [0.003, 0.007]), and a significant interaction effect between motion trajectory and distance at offset (angular with EPS offset vs. radial with PPS offset, *β* = 0.107; *95% CI* [0.02, 0.20]). The effect of distance at offset was nonsignificant (EPS vs. PPS, *β* = 0.062; *95% CI* [– 0.009, 0.13]).

**Figure 3.**
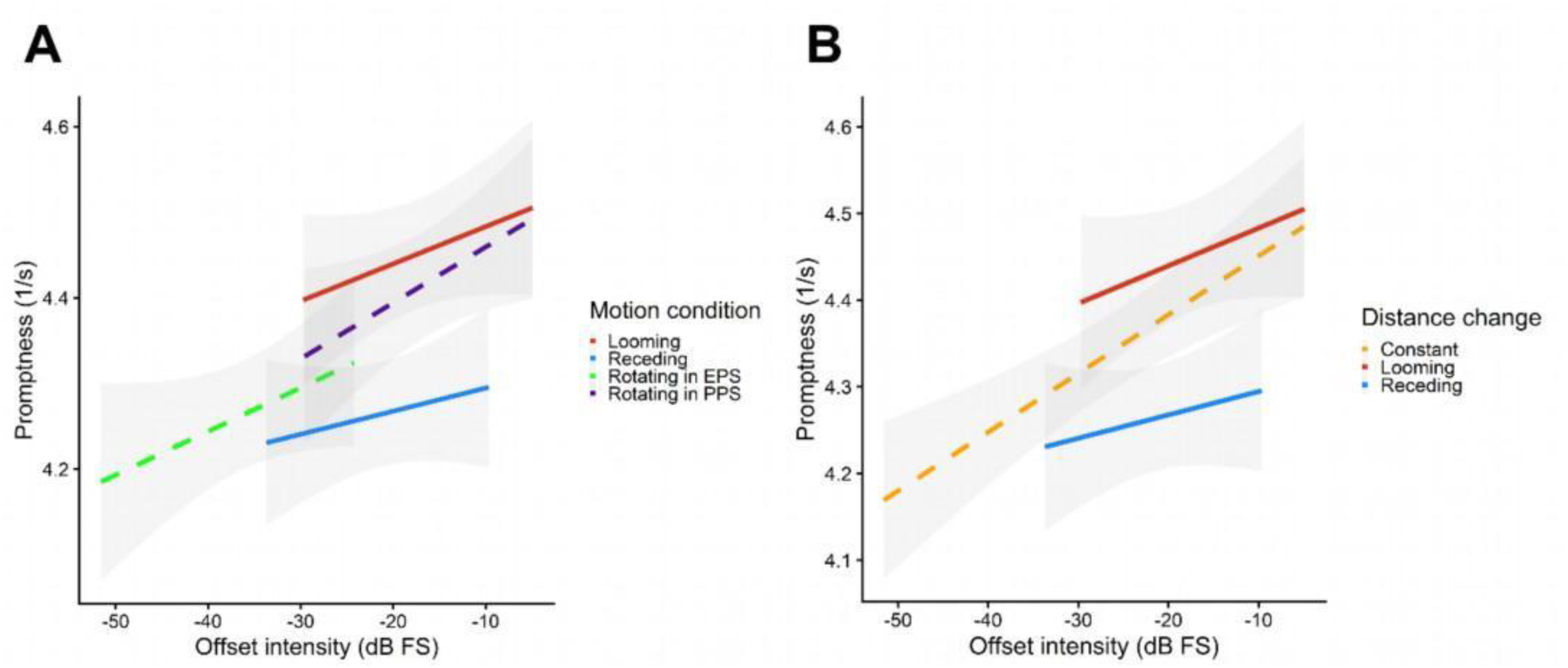
Target detection promptness (1/RT) as a function of offset intensity, motion condition, and source intensity. **A**: Smoothed conditional means of the four motion conditions with standard error (grey sleeves). **B:** Smoothed conditional means based on the winning model, i.e., pooling the two angular conditions (cf. Table 1).

**Table 1.** Model coefficients of the offset distance model and the distance change model predicting promptness. BIC: Bayesian Information Criterion. Smaller BIC values correspond to a more parsimonious model.

|  | Offset distance model | Distance change model |
|---|---|---|
| (Intercept) | 4.435 [3.950, 4.919]*** | 4.492 [4.012, 4.972]*** |
| Intensity at offset | 0.005 [0.003, 0.007]*** | 0.006 [0.004, 0.008]*** |
| Motion trajectory (radial) | -0.070 [-0.136, -0.004]* |  |
| Distance at offset (PPS) | 0.062 [-0.009, 0.133] |  |
| Radial trajectory X PPS offset | 0.107 [0.018, 0.196]* |  |
| Distance change (looming) |  | 0.059 [0.007, 0.112]* |
| Distance change (receding) |  | -0.106 [-0.157, -0.055]*** |
| BIC | 24 784.5 | 24 773.5 |
| ΔBIC | 0 | -11 |

\* p < 0.05, \*\* p < 0.01, \*\*\* p < 0.001

To further clarify the lack of significant effect of offset distance on target detection promptness, we compared the above-mentioned model to one that excluded offset distance as a predictor and instead included a motion direction variable with the factors *looming, receding,* and *constant* (i.e., pooling the two angular conditions). Based on model performance metrics, this model was more parsimonious than the previous model (Δ*BIC* = –11; Table 1). This suggests that disregarding distance in sounds with angular motion improves predictive power, supporting the notion that sounds moving at a near but constant distance do not increase phasic alertness compared to sounds at a far distance on top of the intensity-driven increase. Converting the promptness estimates of the winning model to RT values, we can see that reactions were about 2.89 ms faster (95% *CI =* [0.35, 5.42] ms) after looming cues and about 5.38 ms slower (95% *CI =* [-8.06, –2.76] ms) after receding cues than constant-distance (angular) cues. Besides that, increasing the intensity of the offset accelerated reactions by about 0.3 ms on average (95% *CI =* [0.2, 0.4] ms). In contrast, decreasing the distance of an angular sound can result in a RT change anywhere between –0.46 ms and +6.57 ms (based on the weaker model).

Further, the spatial congruence between cue and target (i.e., whether the target was presented on the same side where the cue ended) and its effect on target detection promptness was investigated in a linear mixed model. Congruence had no main effect on promptness (*M_adj_* = 0.02, *SE* = 0.02, 95% CI [-0.02, 0.06], *p* = 0.41), nor did it interact with either motion condition (looming: *M_adj_* = 0.02, *SE* = 0.04, *95% CI* [-0.07, 0.10], *p* = 0.68; receding: *M_adj_* = 0.06, *SE* = 0.04, *95% CI* [-0.02, 0.15], *p* = 0.13; rotating in PPS: *M_adj_* = 0.01, *SE* = 0.04, *95% CI* [-0.09, 0.07], *p* = 0.83; rotating in EPS: *M_adj_* = –0.002, *SE* = 0.04, *95% CI* [-0.08, 0.08], *p* = 0.96).

### 3.2. EEG results

#### 3.2.1. Interim analysis

In order to determine the point at which data collection should be terminated, two interim analyses (looks) were performed, testing H2 (sound in PPS elicit different responses from sounds in EPS) with a cluster-based permutation analysis on the difference waveform between rotating in EPS and rotating in PPS. At look 1, the sample size was 10, and a nonsignificant effect was found (adjusted alpha = 0.017; smallest p = 0.14; value of the alpha spending function = 0.017). At the second look, the sample size (after exclusions) was 22, and the effect was again nonsignificant (adjusted alpha = 0.02; smallest p = 0.82; value of the alpha spending function = 0.037). Therefore, data collection was terminated at 22 participants (sample size after exclusions as described in the Participants section).

#### 3.2.2. ERP and source localization results

To test whether looming sounds increase sensory gain compared to receding sounds (H1), ERPs were analyzed in the time window of –200 to 1000 ms from motion onset, as this auditory event was expected to evoke an N1-P2 complex based on previous literature (Baumgartner et al., 2017; Getzmann & Lewald, 2010). A cluster-based permutation analysis was run on the looming minus receding difference waveform with 1000 permutations. ERPs evoked by the two radial cues did not significantly differ from each other in the target time window (smallest *p* = 0.61, summed cluster *t =* 9.78).

To explore ERP differences beyond the time window defined in the preregistration, we ran a separate analysis on an extended time window, encompassing the whole duration of the cue as well as the target (up to 2200 ms after cue motion onset). The cluster-based permutation test on the radial difference waveform (looming minus receding) showed a pair of significant clusters, one involving parietal electrodes (*p* < 0.01, summed cluster *t =* –767.65, *d =* –1.30) and the other involving frontal electrodes (*p <* 0.05, summed cluster *t =* 1481.88, *d =* 1.23), both emerging around 1500 ms after cue motion onset and extending to 2200 ms (see Figure 4A for the parietal part of the dipole).

**Figure 4.**
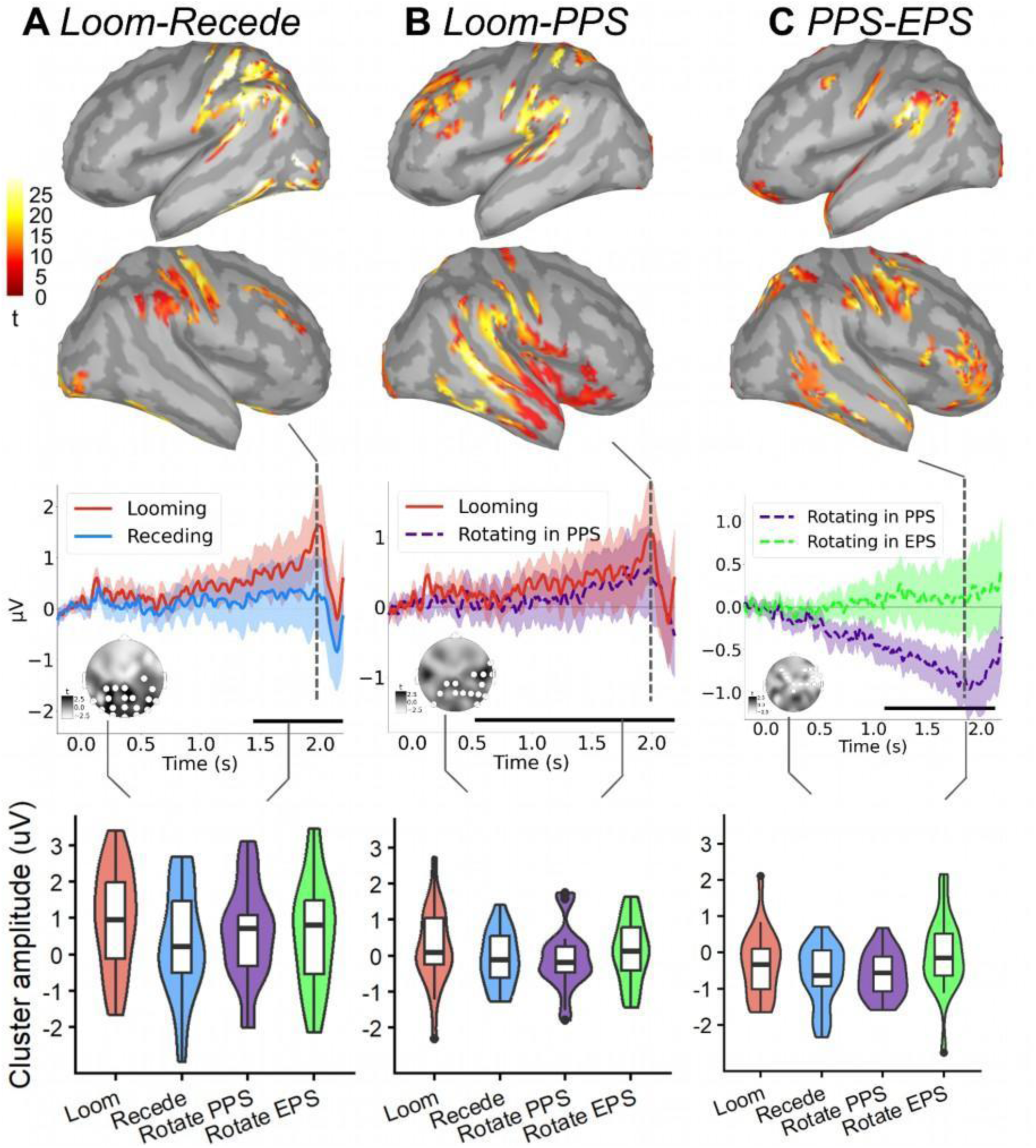
EEG sensor-level clusters and source localization results. **A**: Looming-receding contrast. **B:** Looming-rotating in PPS contrast. **C:** Rotating in EPS – rotating in PPS contrast. **Upper panels:** The activation of the estimated sources of the difference waveforms were tested against the baseline period (–200-0 ms). Significant clusters of source activation are plotted on the brain surface at the time point highlighted in the middle panels with dashed lines (1.98 s after motion onset). The color bars indicate *t*-statistics values. **Middle panels:** ERP time courses measured from cue motion onset at the spatial clusters shown in the lower left corner of each diagram. The time span of a significant cluster is highlighted with a black horizontal line below the waveforms. Colors indicate experimental conditions. **Lower panels:** Cluster amplitude averages in each condition from the spatiotemporal cluster shown on the corresponding column of the middle panel.

The difference between the angular motion conditions (rotating in PPS minus rotating in EPS) was reflected in a central, right-lateralized cluster from around 1500 ms to the end of the epoch at 2200 ms (*p <* 0.05, summed cluster *t =* –1134.74, *d =* –0.93; Supplementary figure 1) – however, when we compared these two conditions in the trials where they had comparable sound intensity (rotating in PPS, low intensity contrasted with rotating in EPS, high intensity), no significant clusters were found, suggesting that the observed difference in cortical processing reflects sound level rather than distance. Finally, the two conditions with PPS offset (looming minus rotating in PPS) were significantly different from each other, with a cluster of parietal electrodes and a temporal extent of about 500 ms to the end of the epoch (*p < 0.05*, summed cluster *t =* 888.29, *d =* 0.99; Figure 4B).

The difference waveforms representing each of the three contrasts were subjected to source localization, and the activation level of the resulting source estimations was tested against the 200 ms baseline period with a parametric Student’s *t*-test. We found differential neural processing of looming and receding sounds predominantly in left hemispheric sensory areas (Figure 4A); sounds ending in PPS after travelling an approaching vs. a constant-distance trajectory in bilateral somatosensory, motor and auditory areas (Figure 4B); and of high and low intensity sounds predominantly in right hemispheric areas (Figure 4C). The specific brain regions involved in each contrast are summarized in Table 2.

**Table 2.** Cortical sources involved in each ERP contrast. L: left hemisphere, R: right hemisphere. The naming convention follows the DKT atlas.

|  | **Looming - Receding** | **Looming - Rotating in PPS** | **Rotating in PPS - Rotating in EPS** |
|---|---|---|---|
| **Frontal** | Superior frontal R<br>Caudal middle frontal R<br>Precentral R | Superior frontal L<br>Middle frontal L<br>Precentral L, R<br>Inferior frontal R | Precentral L, R<br>Middle frontal R<br>Inferior frontal R |
| **Parietal** | Postcentral L, R<br>Supramarginal L<br>Superior parietal L<br>Intraparietal L | Superior parietal L<br>Postcentral L, R<br>Supramarginal L, R<br>Cuneus | Postcentral L, R<br>Supramarginal L, R<br>Precuneus L<br>Superior parietal R |
| **Temporal** | Superior temporal L<br>Middle temporal L<br>Inferior temporal L<br>Fusiform L, R<br>Parahippocampal L, R<br>Lingual L<br>Entorhinal R | Superior temporal L, R<br>Middle temporal R<br>Inferior temporal R<br>Fusiform R | Superior temporal L, R<br>Middle temporal R<br>Inferior temporal R<br>Fusiform R<br>Parahippocampal R<br>Insula R |
| **Occipital** | Lateral occipital L | Medial occipital L, R | - |

### 3.3. SCR results

SCR was analyzed as another indicator of phasic alertness (besides target detection promptness) and was expected to be larger after looming than receding sounds (H1), as well as after sounds ending in PPS rather than in EPS (H2). A cluster-based permutation test with 1000 permutations was run on the SCR data comparing the four conditions (looming, receding, rotating in PPS, rotating in EPS). The initial clustering found no clusters exceeding the significance threshold (alpha = 0.05). Thus, we found no evidence that SCR is differentially affected by different trajectories and distances of auditory motion.

### 3.4. Relationship between neural and behavioral measures

We explored the connection between behavior and cortical signals in the target detection task, i.e., whether ERP amplitudes at the observed looming-receding, looming-rotating in PPS, and rotating in PPS-rotating in EPS difference clusters predict target detection promptness in each trial (Figure 4A-C, lower panels). To this end, we fitted two neurobehavioral models in addition to the behavioral model described in Table 1 (Distance change model) and compared them to find out whether including neural data improved model fit.

In the neurobehavioral models, the same significant offset intensity effects were observed as in the behavioral model, with an additional significant effect of the ERP amplitude in the looming-receding cluster (*β* = 0.008; *95% CI* [0.004, 0.012], *p* < 0.001), in the looming-rotating in PPS cluster (*β* = 0.011; *95% CI* [0.006, 0.017], *p* < 0.001), as well as in the rotating in PPS-rotating in EPS cluster (*β* = 0.006; *95% CI* [0.000, 0.011], *p* < 0.05). Receding distance change had a significant effect on target detection promptness in all 3 neurobehavioral models, but looming distance change did not reach significance when including the looming-receding or the looming-rotating in PPS cluster amplitudes in the model. However, the purely behavioral model was found to be a more parsimonious fit than either of the neurobehavioral ones (Table 3). Thus, although cortical activity at the spatiotemporal clusters underlying the differential processing of radially and angularly moving sounds explains a significant amount of variance in target detection speed, the explained additional variance is not large enough to justify an increase in model complexity (Table 3).

**Table 3.** Model coefficients of the behavioral and neurobehavioral models predicting promptness. BIC: Bayesian Information Criterion. Smaller – in this case more negative – BIC values correspond to a more parsimonious model.

|  | **Behavioral** | **Neurobehavioral (looming-receding cluster)** | **Neurobehavioral (looming-PPS cluster)** | **Neurobehavioral (PPS-EPS cluster)** |
|---|---|---|---|---|
| (Intercept) | 4.536 [4.054, 5.018]*** | 4.531 [4.050, 5.012]*** | 4.534 [4.053, 5.015]*** | 4.538 [4.056, 5.021]*** |
| Intensity at offset (dB FS) | 0.006 [0.004, 0.008]*** | 0.006 [0.004, 0.008]*** | 0.006 [0.004, 0.008]*** | 0.006 [0.004, 0.008]*** |
| Distance change (looming) | 0.063 [0.000, 0.126]* | 0.061 [-0.001, 0.124] | 0.062 [-0.001, 0.124] | 0.063 [0.001, 0.126]* |
| Distance change (receding) | -0.132 [-0.193, -0.071]*** | -0.131 [-0.192, -0.070]*** | -0.132 [-0.192, -0.071]*** | -0.131 [-0.192, -0.070]*** |
| ERP amplitude on Loom-Rec cluster |  | 0.008 [0.004, 0.012]*** |  |  |
| ERP amplitude on Loom-PPS cluster |  |  | 0.011 [0.006, 0.017]*** |  |
| ERP amplitude on PPS-EPS cluster |  |  |  | 0.006 [0.000, 0.011]* |
| BIC | 17 704.3 | 17 709.5 | 17 707.2 | 17 718.7 |
| ΔBIC | 0 | 5.2 | 2.9 | 14.4 |

\* p < 0.05, \*\* p < 0.01, \*\*\* p < 0.001

## 4. Discussion

In this study, we tested the manifestation of the auditory looming bias (ALB) and the auditory peripersonal space (PPS) bias and explored their relationship. We used an auditory target detection task with sound sources moving at different distances and trajectories and additionally assessed distance perception in stationary and moving sources relative to the listener’s body. We found evidence of the ALB affecting phasic alertness and sensory gain, but we could not confirm that there is a more general perceptual bias towards sources within PPS. Additionally, we found an overall tendency to perceive moving sounds as approaching.

### 4.1. Looming auditory motion increases phasic alertness and sensory gain

We found increased target detection promptness and ERP amplitudes following looming sounds compared to both receding sounds and sounds moving at a constant distance. The difference was reflected in cortical areas attributed to multiple sensory systems and their integration. These results underline the preferential processing of looming auditory motion, corroborating H1.

The increased alerting response to looming as opposed to receding auditory motion is in line with previous studies. Looming sounds accelerate subsequent target detection whether the approach is signaled by intensity (Bach et al., 2008, 2015) or full motion cues (Bach et al., 2009), while audiovisual motion detection speed also increases if the motion is looming rather than receding (Cappe et al., 2012). Thus, looming sounds seem to increase alerting-related behavioral facilitation and response readiness.

Further, the increased sensory gain during the neural processing of looming sounds sources also agrees with previous findings of larger sustained cortical activity while tracking the behavioral relevance of looming as opposed to receding sounds (Bach et al., 2015). Other studies found transient rather than sustained cortical differentiation of looming and receding motion (Baumgartner et al., 2017; Greif et al., 2025; Ignatiadis et al., 2024). Methodological differences between the two groups of studies suggest that transient responses reflect an attentional focus on the motion onset, while sustained activity underlies the preparation for a subsequent event. Both of these processes are distinctly affected by looming and receding motion.

The estimated cortical generators of the sustained scalp activity in right-hemispheric frontal areas are implicated in the alerting network (Posner et al., 2006) and are compatible with previous studies showing motor and premotor activation in the auditory looming bias (Ignatiadis et al., 2025; Seifritz et al., 2002). Not only the looming—receding, but also the looming—near rotation contrast was characterized by activity in motor preparation-related areas, while targets were detected faster after looming sounds compared to all control conditions. Looming sonic motion thus seems to increase motor preparation not only compared to motion in the opposite direction, but even relative to motion near the body.

Additionally, multisensory integration areas were differentially activated by looming sounds compared to both receding and near sounds. Differences were found in auditory, somatosensory, and visual areas, as well as regions implicated in spatial processing. Previous studies also found activation in visual areas evoked by looming sounds (Bach et al., 2015; Seifritz et al., 2002), suggesting that the association of looming auditory stimuli with other sensory systems might be particularly strong. The activation of the inferior parietal lobule, a center of multisensory integration (Huang et al., 2005; Renier et al., 2009; Sun et al., 2022), further suggests that multiple sensory systems were recruited, increasing the likelihood of detecting potential objects in the visual field or on the skin. Thus, our results are compatible with the notion that approaching objects increase multisensory integration (Cappe et al., 2012; Cléry et al., 2015, 2017; Maier et al., 2004) compared to receding and near but constant-distance motion. Note however that activation in a wide range of cortical areas during a unisensory task is not direct evidence of multisensory integration, and our source estimation results are likely affected by source spread and inverse solution uncertainty.

Further looming—receding differences emerged in regions commonly exhibiting repetition suppression, i.e., a reduction of activity in response to repeated stimuli (left ventral temporal cortex; Buckner et al., 2000; Fiebach et al., 2005; Lee et al., 2020; McDonald et al., 2010). Affective stimuli lead to decreased repetition suppression (for a review see Trapp & Kotz, 2016), while looming sounds are more affectively loaded (Bach et al., 2009). Because repetition probability was equal across motion trajectories in our experiment, this suggests that looming sounds might be less susceptible to repetition suppression than receding sounds. Future studies could directly test this interpretation.

Finally, cortical activation differences specific to the looming—near rotation contrast coincided with the dorsal auditory stream for sound localization (van der Heijden et al., 2019). This might reflect the role of localizability in the ALB (Greif et al., 2025). Passive listening tasks with spatialized sounds evoke activity in the hemisphere contralateral to sound presentation (van der Heijden et al., 2019), so the bilateral activation observed here likely stems from the presentation of sound on both sides, rather than a strong integration of information from both auditory hemifields. Overall, looming sounds seem to increase the activation of the alerting network not only relative to receding (Canzoneri et al., 2012; Finisguerra et al., 2015) and stationary (Camponogara et al., 2015) sounds but also compared to sounds moving within the immediate vicinity of the body.

### 4.2. No evidence for a general bias towards sounds within PPS

H2 proposed a bias toward sound sources in PPS which increases the effect of the ALB. We did not find evidence to support this hypothesis. Alertness and sensory gain did not increase after sounds ending in PPS in general, only if the motion trajectory was looming. Moreover, the offset distance of angular sources could not explain any variance in target detection promptness over and above intensity differences. Thus, we suggest that the ALB may not be further amplified by a preferential processing of sound sources reaching the immediate environment of the body.

It is important to acknowledge that sounds simulated in EPS were still largely perceived within arm’s reach according to the distance rating task; i.e., based on the current stimuli, auditory PPS – at least as defined by the portion of space where the arm can act – was not clearly demarcated by our participants. Nevertheless, sounds simulated in EPS were perceived farther (at around arm’s length) than those simulated in PPS; thus, relatively near and far sound sources were perceived.

In sounds with angular motion, distance did not explain phasic alertness effects over and above sound intensity. Rather, sounds ending near the body only affected phasic alertness following an approaching trajectory. Although this is puzzling in light of previous reports of objects in PPS increasing alertness, our results are in line with previous studies upon closer inspection. Namely, findings about motor preparation scaling with decreasing distance were specific to looming sounds, absent for near stationary sounds (Bahadori et al., 2021; Camponogara et al., 2015), and weaker for the near portion of receding sounds (Canzoneri et al., 2012). Thus, our results agree with the previous literature in that the role of near auditory distance in increasing alertness is specific to approaching sources.

In contrast, Finisguerra and colleagues (2015) reported higher motor excitability for sounds in PPS regardless of looming or receding direction. In the current study, we observed activation differences near the motor cortex in the looming—receding and looming—near rotation contrasts. However, the latencies are quite different: 50 ms in Finisguerra and colleagues’ study compared to hundreds of milliseconds in our results and other studies (Bahadori et al., 2021; Camponogara et al., 2015; Canzoneri et al., 2012). Indeed, motor excitability in response to near sounds decreases with time (Serino et al., 2009). Taken these effects together, an auditory PPS bias might govern early corticospinal responses but not affect alertness states emerging in later processing stages unless its effects are combined with the ALB.

We also found no evidence that sounds presented close to the body increase overall sensory gain independently of the acoustic intensity, unless, again, the sound source approaches from a distance. Looming sounds increased neural responses more than sounds moving at a constant distance, reflected in left-hemispheric motor cortical areas, as well as somatosensory and auditory areas bilaterally.

### 4.3. Sound offsets are perceived as being closer than sound onsets

Interestingly, listeners tended to perceive sound offsets closer than sound onsets, even if the actual distance was constant. Previous studies found that the offset of looming sounds is perceived closer than the equidistant offset of receding sounds (Neuhoff, 2001; Neuhoff et al., 2009), reflecting a perceptual bias towards the direction of displacement, termed representational momentum (Hubbard, 2014). However, representational momentum cannot fully account for the current findings that sound sources changing only in azimuth are also perceived to end closer than they started. At the same time, the offsets of moving sounds were perceived closer than the overall distance of stationary sounds – which is already largely underestimated (Kolarik et al., 2016). Therefore, our results hint at a perceptual bias governing the distance perception of moving sound offsets specifically. Alternatively, the effect can be explained by memory processes, as sound offsets are more recent and less masked by a relatively long sound than sound onsets. Even so, the direction of the effect, namely that offsets were rated as closer rather than farther than onsets, might reflect a defensive bias. However, because our distance rating task was not originally designed to test this idea, more systematic investigations are needed, for example by collecting separate onset and offset responses to stationary sounds as well.

## 5. Conclusion

It is debated whether near auditory sources are processed preferentially compared to far sources, independently of their motion direction. We addressed this question by examining distance perception and target detection performance related to sounds moving at different distances and directions. In terms of distance perception, we observed a tendency to perceive sound offsets as closer than onsets in moving sounds. Additionally, since our results also suggest that auditory distance perception might be different between moving and stationary sources, we encourage the use of moving but constant-distance controls in the study of the auditory PPS and auditory distance perception.

We further found that looming auditory motion increases phasic alertness not only compared to receding sounds but also compared to sounds moving near the body at a constant distance. We did not find evidence that auditory motion near the body increases alertness over and above the effects of sound intensity. These conclusions are compatible with the relative imprecision of auditory distance recognition and are also in line with broader models of perception that emphasize change detection over state detection.

## Acknowledgment

This work was supported by the European Union’s Horizon 2020 framework program for research and innovation under the Marie Skłodowska-Curie Grant Agreement No: 101120054 to PK, RB and FP.

## Data availability

Data can be accessed at https://doi.org/10.60887/n2tamqt6r3pk8bk7. Experimental and analysis scripts are available on GitHub (https://github.com/kovacs-petra/cherish).

## Statement on the use of generative AI

Generative AI (GPT-5) was used for coding tasks and to make the final manuscript more concise. The authors reviewed and edited the suggested changes and take full responsibility for the manuscript and related codes.

## References

1. Armstrong, C., Thresh, L., Murphy, D., & Kearney, G. (2018). A perceptual evaluation of individual and non-individual HRTFs: A case study of the SADIE II database. Applied Sciences, 8(11), 2029.

2. Bach, D. R., Furl, N., Barnes, G., & Dolan, R. J. (2015). Sustained magnetic responses in temporal cortex reflect instantaneous significance of approaching and receding sounds. PLoS One, 10(7), e0134060. 10.1371/journal.pone.0134060

3. Bach, D. R., Neuhoff, J. G., Perrig, W., & Seifritz, E. (2009). Looming sounds as warning signals: The function of motion cues. International Journal of Psychophysiology, 74(1), 28–33. 10.1016/j.ijpsycho.2009.06.004

4. Bach, D. R., Schachinger, H., Neuhoff, J. G., Esposito, F., Salle, F. D., Lehmann, C., Herdener, M., Scheffler, K., & Seifritz, E. (2008). Rising Sound Intensity: An Intrinsic Warning Cue Activating the Amygdala. Cerebral Cortex, 18(1), 145–150. 10.1093/cercor/bhm040

5. Bahadori, M., Barumerli, R., Geronazzo, M., & Cesari, P. (2021). Action planning and affective states within the auditory peripersonal space in normal hearing and cochlear-implanted listeners. Neuropsychologia, 155, 107790. 10.1016/j.neuropsychologia.2021.107790

6. Bates, D., Mächler, M., Bolker, B., & Walker, S. (2015). Fitting linear mixed-effects models using lme4. Journal of Statistical Software, 67, 1–48. 10.18637/jss.v067.i01

7. Baumgartner, R., Reed, D. K., Tóth, B., Best, V., Majdak, P., Colburn, H. S., & Shinn-Cunningham, B. (2017). Asymmetries in behavioral and neural responses to spectral cues demonstrate the generality of auditory looming bias. Proceedings of the National Academy of Sciences, 114(36), 9743–9748. 10.1073/pnas.1703247114

8. Brungart, D. S., Durlach, N. I., & Rabinowitz, W. M. (1999). Auditory localization of nearby sources. II. Localization of a broadband source. The Journal of the Acoustical Society of America, 106(4), 1956–1968.

9. Buckner, R. L., Koutstaal, W., Schacter, D. L., & Rosen, B. R. (2000). Functional MRI evidence for a role of frontal and inferior temporal cortex in amodal components of priming. Brain, 123(3), 620–640.

10. Camponogara, I., Komeilipoor, N., & Cesari, P. (2015). When distance matters: Perceptual bias and behavioral response for approaching sounds in peripersonal and extrapersonal space. Neuroscience, 304, 101–108.

11. Canzoneri, E., Magosso, E., & Serino, A. (2012). Dynamic sounds capture the boundaries of peripersonal space representation in humans. PLoS One, 7(9).

12. Cappe, C., Thelen, A., Romei, V., Thut, G., & Murray, M. M. (2012). Looming signals reveal synergistic principles of multisensory integration. Journal of Neuroscience, 32(4), 1171– 1182.

13. Cléry, J., Guipponi, O., Odouard, S., Pinède, S., Wardak, C., & Ben Hamed, S. (2017). The Prediction of Impact of a Looming Stimulus onto the Body Is Subserved by Multisensory Integration Mechanisms. The Journal of Neuroscience, 37(44), 10656–10670. 10.1523/JNEUROSCI.0610-17.2017

14. Cléry, J., Guipponi, O., Odouard, S., Wardak, C., & Hamed, S. B. (2015). Impact prediction by looming visual stimuli enhances tactile detection. Journal of Neuroscience, 35(10), 4179–4189.

15. Cuevas-Rodríguez, M., Picinali, L., González-Toledo, D., Garre, C., de la Rubia-Cuestas, E., Molina-Tanco, L., & Reyes-Lecuona, A. (2019). 3D Tune-In Toolkit: An open-source library for real-time binaural spatialisation. PloS One, 14(3), e0211899.

16. Delorme, A., & Makeig, S. (2004). EEGLAB: an open source toolbox for analysis of single-trial EEG dynamics including independent component analysis. Journal of Neuroscience Methods, 134(1), 9–21.

17. Fiebach, C. J., Gruber, T., & Supp, G. G. (2005). Neuronal mechanisms of repetition priming in occipitotemporal cortex: Spatiotemporal evidence from functional magnetic resonance imaging and electroencephalography. Journal of Neuroscience, 25(13), 3414–3422.

18. Finisguerra, A., Canzoneri, E., Serino, A., Pozzo, T., & Bassolino, M. (2015). Moving sounds within the peripersonal space modulate the motor system. Neuropsychologia, 70, 421– 428.

19. Getzmann, S., & Lewald, J. (2010). Effects of natural versus artificial spatial cues on electrophysiological correlates of auditory motion. Hearing Research, 259(1–2), 44–54.

20. Ghazanfar, A. A., Neuhoff, J. G., & Logothetis, N. K. (2002). Auditory looming perception in rhesus monkeys. Proceedings of the National Academy of Sciences, 99(24), 15755– 15757.

21. González-Toledo, D., Molina-Tanco, L., Cuevas-Rodríguez, M., Majdak, P., & Reyes-Lecuona, A. (2023). The Binaural Rendering Toolbox. A virtual laboratory for reproducible research in psychoacoustics. Forum Acusticum.

22. Gramfort, A., Luessi, M., Larson, E., Engemann, D. A., Strohmeier, D., Brodbeck, C., Goj, R., Jas, M., Brooks, T., Parkkonen, L., & Hämäläinen, M. S. (2013). MEG and EEG Data Analysis with MNE-Python. Frontiers in Neuroscience, 7(267), 1–13. 10.3389/fnins.2013.00267

23. Graziano, M. S., Reiss, L. A., & Gross, C. G. (1999). A neuronal representation of the location of nearby sounds. Nature, 397(6718), 428–430.

24. Greif, T., Barumerli, R., Ignatiadis, K., Tóth, B., & Baumgartner, R. (2025). The role of spatial perception in auditory looming bias: Neurobehavioral evidence from impossible ears. Frontiers in Neuroscience, 19, 1645936.

25. Hillyard, S. A., Vogel, E. K., & Luck, S. J. (1998). Sensory gain control (amplification) as a mechanism of selective attention: Electrophysiological and neuroimaging evidence. Philosophical Transactions of the Royal Society of London. Series B: Biological Sciences, 353(1373), 1257–1270.

26. Huang, M.-X., Lee, R. R., Miller, G. A., Thoma, R. J., Hanlon, F. M., Paulson, K. M., Martin, K., Harrington, D. L., Weisend, M. P., Edgar, J. C., & others. (2005). A parietal–frontal network studied by somatosensory oddball MEG responses, and its cross-modal consistency. Neuroimage, 28(1), 99–114.

27. Hubbard, T. L. (2014). Forms of momentum across space: Representational, operational, and attentional. Psychonomic Bulletin & Review, 21(6), 1371–1403.

28. Ignatiadis, K., Baier, D., Barumerli, R., Sziller, I., Tóth, B., & Baumgartner, R. (2024). Cortical signatures of auditory looming bias show cue-specific adaptation between newborns and young adults. Communications Psychology, 2(1), 56.

29. Ignatiadis, K., Barumerli, R., Deco, G., Tóth, B., & Baumgartner, R. (2025). Threat-related corticocortical connectivity elicited by rapid auditory looms. Scientific Reports.

30. Kolarik, A. J., Moore, B. C., Zahorik, P., Cirstea, S., & Pardhan, S. (2016). Auditory distance perception in humans: A review of cues, development, neuronal bases, and effects of sensory loss. *Attention, Perception*, & Psychophysics, 78(2), 373–395. 10.3758/s13414-015-1015-1

31. Kopčo, N., Huang, S., Belliveau, J. W., Raij, T., Tengshe, C., & Ahveninen, J. (2012). Neuronal representations of distance in human auditory cortex. Proceedings of the National Academy of Sciences, 109(27), 11019–11024. 10.1073/pnas.1119496109

32. Kopčo, N., & Shinn-Cunningham, B. G. (2011). Effect of stimulus spectrum on distance perception for nearby sources. The Journal of the Acoustical Society of America, 130(3), 1530–1541.

33. Kovács, P., Winkler, I., Tóth, B., Pavani, F., & Baumgartner, R. (2025). Auditory peripersonal space and the looming bias in normal hearing adults. OSF Registries. 10.17605/OSF.IO/MB2GA

34. Ladavas, E., Pavani, F., & Farne, A. (2001). Auditory peripersonal space in humans: A case of auditory-tactile extinction. Neurocase, 7(2), 97–103.

35. Lee, S.-M., Henson, R. N., & Lin, C.-Y. (2020). Neural correlates of repetition priming: A coordinate-based meta-analysis of fMRI studies. Frontiers in Human Neuroscience, 14, 565114.

36. Lenth, R. (2023). emmeans: Estimated Marginal Means, aka Least-Squares Means_. R Package Version 1.8. 5.

37. Little, A. D., Mershon, D. H., & Cox, P. H. (1992). Spectral Content as a Cue to Perceived Auditory Distance. Perception, 21(3), 405–416. 10.1068/p210405

38. Maier, J. X., Neuhoff, J. G., Logothetis, N. K., & Ghazanfar, A. A. (2004). Multisensory integration of looming signals by rhesus monkeys. Neuron, 43(2), 177–181.

39. Majdak, P., Brinkmann, F., De Muynke, J., Mihocic, M., & Noisternig, M. (2022). Spatially oriented format for acoustics 2.1: Introduction and recent advances. Journal of the Audio Engineering Society, 70, 565–584.

40. McDonald, C. R., Thesen, T., Carlson, C., Blumberg, M., Girard, H. M., Trongnetrpunya, A., Sherfey, J. S., Devinsky, O., Kuzniecky, R., Dolye, W. K., & others. (2010). Multimodal imaging of repetition priming: Using fMRI, MEG, and intracranial EEG to reveal spatiotemporal profiles of word processing. Neuroimage, 53(2), 707–717.

41. Mehrpour, V., Martinez-Trujillo, J. C., & Treue, S. (2020). Attention amplifies neural representations of changes in sensory input at the expense of perceptual accuracy. Nature Communications, 11(1), 2128. 10.1038/s41467-020-15989-0

42. Moore, B. C., Jervis, M., Harries, L., & Schlittenlacher, J. (2018). Testing and refining a loudness model for time-varying sounds incorporating binaural inhibition. The Journal of the Acoustical Society of America, 143(3), 1504–1513.

43. Neuhoff, J. G. (1998). Perceptual bias for rising tones. Nature, 395(6698), 123–124.

44. Neuhoff, J. G. (2001). An adaptive bias in the perception of looming auditory motion. Ecological Psychology, 13(2), 87–110.

45. Neuhoff, J. G. (2016). Looming sounds are perceived as faster than receding sounds. Cognitive Research: Principles and Implications, 1(1), 15.

46. Neuhoff, J. G., Long, K. L., & Worthington, R. C. (2012). Strength and physical fitness predict the perception of looming sounds. Evolution and Human Behavior, 33(4), 318–322. 10.1016/j.evolhumbehav.2011.11.001

47. Oostenveld, R., Fries, P., Maris, E., & Schoffelen, J.-M. (2011). FieldTrip: Open Source Software for Advanced Analysis of MEG, EEG, and Invasive Electrophysiological Data. Computational Intelligence and Neuroscience, 2011, 1–9. 10.1155/2011/156869

48. Pascual-Marqui, R. D. (2002). Standardized low-resolution brain electromagnetic tomography (sLORETA): Technical details. Methods Find Exp Clin Pharmacol, 24(Suppl D), 5–12.

49. Petersen, S. E., & Posner, M. I. (2012). The Attention System of the Human Brain: 20 Years After. Annual Review of Neuroscience, 35(1), 73–89. 10.1146/annurev-neuro-062111-150525

50. Posner, M. I., Sheese, B. E., Odludaş, Y., & Tang, Y. (2006). Analyzing and shaping human attentional networks. Neural Networks, 19(9), 1422–1429.

51. R Core Team. (2024). R: A Language and Environment for Statistical Computing. R Foundation for Statistical Computing. https://www.R-project.org/

52. Renier, L. A., Anurova, I., De Volder, A. G., Carlson, S., VanMeter, J., & Rauschecker, J. P. (2009). Multisensory integration of sounds and vibrotactile stimuli in processing streams for “what” and “where.” Journal of Neuroscience, 29(35), 10950–10960.

53. Rosenblum, L. D., Carello, C., & Pastore, R. E. (1987). Relative Effectiveness of Three Stimulus Variables for Locating a Moving Sound Source. Perception, 16(2), 175–186. 10.1068/p160175

54. Rosenblum, L. D., Wuestefeld, A. P., & Saldaña, H. M. (1993). Auditory Looming Perception: Influences on Anticipatory Judgments. Perception, 22(12), 1467–1482. 10.1068/p221467

55. Seifritz, E., Neuhoff, J. G., Bilecen, D., Scheffler, K., Mustovic, H., Schächinger, H., Elefante, R., & Di Salle, F. (2002). Neural Processing of Auditory Looming in the Human Brain. Current Biology, 12(24), 2147–2151. 10.1016/S0960-9822(02)01356-8

56. Serino, A., Annella, L., & Avenanti, A. (2009). Motor properties of peripersonal space in humans. PloS One, 4(8), e6582.

57. Sun, J., Huang, J., Wang, A., Zhang, M., & Tang, X. (2022). The role of the interaction between the inferior parietal lobule and superior temporal gyrus in the multisensory Go/No-go task. NeuroImage, 254, 119140.

58. Tadel, F., Baillet, S., Mosher, J. C., Pantazis, D., & Leahy, R. M. (2011). Brainstorm: A user-friendly application for MEG/EEG analysis. Computational Intelligence and Neuroscience, 2011(1), 879716.

59. The MathWorks Inc. (2023). MATLAB version: 23.2.0 (R2023b) [Computer software]. The MathWorks Inc. https://www.mathworks.com

60. Trapp, S., & Kotz, S. A. (2016). Predicting affective information–An evaluation of repetition suppression effects. Frontiers in Psychology, 7, 1365.

61. van der Heijden, K., Rauschecker, J. P., de Gelder, B., & Formisano, E. (2019). Cortical mechanisms of spatial hearing. Nature Reviews Neuroscience, 20(10), 609–623.

62. Veillette, J. P., Ho, L., & Nusbaum, H. C. (2023). Permutation-based group sequential analyses for cognitive neuroscience. Neuroimage, 277, 120232.

63. Zahorik, P., & Wightman, F. L. (2001). Loudness constancy with varying sound source distance. Nature Neuroscience, 4(1), 78–83. 10.1038/82931

64. Zampini, M., Torresan, D., Spence, C., & Murray, M. M. (2007). Auditory–somatosensory multisensory interactions in front and rear space. Neuropsychologia, 45(8), 1869–1877.

65. Zanini, A., Patané, I., Blini, E., Salemme, R., Koun, E., Farnè, A., & Brozzoli, C. (2021). Peripersonal and reaching space differ: Evidence from their spatial extent and multisensory facilitation pattern. Psychonomic Bulletin & Review, 28(6), 1894–1905.

